# Fast multi-directional DSLM for confocal detection without striping artifacts

**DOI:** 10.1101/2020.04.06.027037

**Authors:** Pietro Ricci, Giuseppe Sancataldo, Vladislav Gavryusev, Alessandra Franceschini, Marie Caroline Müllenbroich, Ludovico Silvestri, Francesco Saverio Pavone

**Affiliations:** European Laboratory for Non-Linear Spectroscopy, Sesto Fiorentino, 50019, Italy; University of Florence, Department of Physics and Astronomy, Sesto Fiorentino, 50019, Italy; University of Palermo, Department of Physics and Chemistry, Palermo, 90128, Italy; National Institute of Optics, Sesto Fiorentino, 50019, Italy; School of Physics and Astronomy, University of Glasgow, G12 8QQ, Glasgow, UK

## Abstract

In recent years light-sheet fluorescence microscopy (LSFM) has become a cornerstone technology for neuroscience, improving quality and capabilities of 3D imaging. By selectively illuminating a single plane, it provides intrinsic optical sectioning and fast image recording, while minimizing out of focus fluorescence background, sample photo-damage and photo-bleaching. However, images acquired with LSFM are often affected by light absorption or scattering effects, leading to un-even illumination and striping artifacts. In this work we present an optical solution to this problem, via fast multi-directional illumination of the sample, based on an acousto-optical deflector (AOD). We demonstrate that this pivoting system is compatible with confocal detection in digital scanned laser light-sheet fluorescence microscopy (DSLM) by using a pivoted elliptical-Gaussian beam. We tested its performance by acquiring signals emitted by specific fluorophores in several mouse brain areas, comparing the pivoting beam illumination and a traditional static one, measuring the point spread function response and quantifying the striping reduction. We observed real-time shadow suppression, while preserving the advantages of confocal detection for image contrast.

## 1. Introduction

Light-sheet fluorescence microscopy (LSFM) is rapidly becoming a landmark for neuroscience and biological processes visualization [1, 2]. It allows 3D imaging of biological samples with high frame rate and micron-scale spatial resolution. Thanks to these properties, LSFM has been exploited in a wide rage of applications from live imaging of fast processes up to long-term tracking of biological dynamics [3–8].

A thin light-sheet, usually created by a cylindrical lens or by rapidly sweeping a collimated beam (as in digital scanned laser light-sheet fluorescence microscope - DSLM), is projected and scanned through the sample, optically sectioning the volume plane by plane. The optical system provides the illumination of a single plane, while a wide-field detection apparatus records, in the orthogonal direction, the fluorescence signal emitted by specific fluorophores [9]. Out-of-focus contributions are avoided by exciting only the focal plane inside the sample. With such intrinsic optical sectioning capability, fluorescence background is minimized, together with sample photo-damage and photo-bleaching.

Both thin and extended samples have been studied over the last years with LSFM, monitoring embryo development or exploring large nervous areas of flies, fishes and small mammals [10–14], up to obtaining whole brain reconstructions with sub-cellular resolution. Thick samples require to be chemically treated before volumetric imaging because tissues are often made of several proteic and lipidic components with different refractive indexes which affect heterogeneously the light interaction. Several optical clearing processes have been introduced and applied to these tissues in order to reduce the refractive index mismatch within the sample [15, 16]. However, residual inhomogeneities lead to artifacts and aberrations in LFSM images, due to light-matter interactions such as scattering or absorption phenomena. Scattered illumination light reduces the optical sectioning capability of LSFM, while scattered emitted photons represent also a detection issue because their trajectories and intensity contribution cannot be distinguished from that of ballistic ones, consequently reducing the image signal-to-noise ratio (SNR) and contrast. Confocal detection (CLSFM) has been introduced to overcome these problems [17, 18]. By synchronizing the rolling shutter of a scientific complementary metal-oxide-semiconductor detector array (sCMOS) - a digital slit made of a few rows of pixels - with the sweep of the illuminating beam, it allows to reject out-of-focus and scattered light.

On the other hand, illuminating samples containing absorbing objects (impurities, blood vessels, pigmentation spots or small air bubbles) produces shadows behind the obstructions - usually named striping artifacts - that severely affect the image quality. Several approaches have been developed to deal with those stripes [19]. Some are based on image post-processing, such as the correction of the artifacts using the information provided by a voxel map of attenuation obtained by a projection tomography over the sample [20]. Other methods act directly on the beam distribution profile used for sample illumination: for instance Bessel beams allowed to solve the striping problems in non-homogeneous media [21–26]. These beams take advantage of their self-healing capability, that is the reconstruction of the original intensity distribution after an obstacle. However, Bessel’s light distribution, which is defined by a typical outer rings structure, carries out-of-focus illumination contributes and leads to image contrast and signal-to-noise ratio degradation with respect to Gaussian beam illumination. A reliable, but complex and time consuming alternative is represented by a multi-view acquisition approach, which requires multi-directional detection and overlapping in post-processing the images taken at different angles [27].

Recently, a new way to address the striping issue has been presented by [28] that enabled a 20 % striping reduction in respect to a classic DSLM. Such improvement has been achieved by inducing small angle axial oscillations (i.e. along the objective detection direction) of the illumination Gaussian beam, realizing, thus, an axially dithered digital scanned light-sheet microscope (aDSLM). A different optical approach is the one proposed by [29] with their diffuse light-sheet microscope. Through the addition of a simple optical element such as a line diffuser, the light propagation through the sample results randomized, allowing the light-sheets to restore after obstructions. Even if an analysis in terms of resolution and contrast compared with other well-known approaches is still missing, it has been demonstrated that striping artifacts were severely reduced, both in a selective plane illumination microscope (SPIM) and in a DSLM configuration. However, such approach is hardly adaptable to the confocal detection modality, because diffusing the light beam breaks the requirement of having a single illumination line sweeping synchronously with the digital slit.

One further method to solve the striping problem leverages a multi-directional illumination approach, where the sample is illuminated from one or two opposite directions by a beam pivoting relative to the focal plane [30]. To realize such multi-directional selective plane illumination microscopy, the beam must rotate faster than the image acquisition rate, i.e. the integration time of the detector, to be able to average out over time the shadow attenuation at different angles. Light-sheet pivoting is usually realized using galvanometric mirrors [30], but they are severely limited by their intrinsic mechanical inertia (peak sweeping rate of 200 Hz for closed-loop mirrors and 12 kHz for resonant ones). For this reason, they are not optimal for a confocal detection regime where, with advanced sCMOS sensors, the line exposure times in rolling shutter modality can be lower than 100 μs, corresponding to sweeping rates in excess of 10 kHz. Furthermore, it has been demonstrated [31] that image artifacts are still evident using slow sweeping rates, while they are greatly reduced at higher rates. The pivoting dynamic can be sped up using acousto-optic deflectors (AODs) which do not have any inertial restrictions [32] and allow to generate simultaneously multiple beams at different angles (static multi-angle illumination) or to rapidly sweep a single beam (dynamic pivoting), reaching up to MHz rates [31]. Moreover, AODs can create and sweep two illumination beams in order to leverage the dual rolling shutter mode of several sCMOS cameras [33]. In this regard, a key alternative has been developed by [34], where the advantages of a multi-directional digital scanned light-sheet microscopy (mDSLM) are merged with the confocal line detection technique. By introducing a cylindrical telescope, an illumination elliptical-Gaussian beam is generated and exploited for striping mitigation without requiring any pivoting, due to the intrinsic degree of “angular diversity” of such profile, although with decreased axial light uniformity that requires image tiling to cover the field of view (FOV) [35].

Here we propose a hybrid DSLM method where each scanning line projected onto the sample is swept by a closed-loop galvo mirror and pivoted around the detection plane by means of an AOD. There is an inherent conflict between the illumination geometry and the finite size of the digital slit used in confocal detection, because a large part of the pivoting beam would rotate out of such digital slit. In the following, we present a method to overcome this issue, while preserving the contrast improvement due to confocal detection, the axial illumination uniformity and simultaneously attenuating striping artifacts. Building upon the work in [34], we exploited the features of two cylindrical lenses to create an optimized optical beam-shaping system that generates an expanded elliptical-Gaussian beam that covers the digital slit while being pivoted by an AOD. We imaged several mouse brain areas, observing real-time shadow suppression while preserving confocal detection of the signal emitted by specific fluorophores and no decrease in axial light uniformity. A comparison between such scanning beam illumination and a standard static one has been carried out in terms of shadowing reduction and point spread function (PSF) response.

## 2. Methods

A custom-made light-sheet fluorescence microscope has been used for whole mouse brains imaging. Its complete detailed description can be found in [36], while here we present the modifications introduced to suppress the striping artifacts.

### 2.1. Multi-Beam and Scan-Beam Light-sheet Fluorescence Microscope

The setup is schematically represented in Figure (1). In respect to the configuration described in [36], we modified one of the two identical excitation arms, introducing an alternative optical path, as indicated with the red dashed rectangle. This pathway is selectable through a pair of flip-flop mirrors placed between the laser unit and the galvanometric scanning system. In detail, the visible light beam generated by a laser diode (Cobolt AB, Jive 561 nm, 50 mW, s-polarized) is expanded 5× by a telescope (Thorlabs AC080-020-A *f_L_* = 20 mm and Thorlabs AC254-200-A *f_L_* = 200 mm) and is then guided through a second telescope made by two cylindrical lenses (Thorlabs ACY254-150-B *f_L_* = 150 mm and Thorlabs LJ1878L2-A *f_L_* = 10 mm) to collapse one beam dimension by a factor of 15. Afterwards, the light goes through a half-wave plate and enters into an AOD (AA Opto Electronic, DTSX-400, TeO_2_, aperture 7.5 × 7.5 mm^2^) which is driven by a RF multi-channel driver (MDSnC, 8 channels, centered at 92 MHz, bandwidth 56 MHz). The pivoted beam is then collected and collimated by a Thorlabs AC508-500-A *f_L_* = 500 mm, which forms a 1:1 telescope with the first lens *f_L_* = 500 mm placed after the second flip-flop mirror. The following galvo head digitally generates the light-sheet and is positioned in a conjugated plane with the back focal plane of the excitation objective Plan Flour EPI, 10x, 0.3NA, WD 17.5 mm, Nikon, Japan. An Olympus XLPlan N 10x/0.60 SVMP Objective, paired with a *f_L_* = 200 mm tube lens, is used to detect the fluorescence emitted by the samples. The signal is then filtered by a bandpass filter (FF01-609/54-25, Semrock) and, finally, collected by a sCMOS camera (Orca Flash4.0 v2.0, Hamamatsu) with a pixel size of 6.5 × 6.5 μm^2^ and an active area of 13.3 × 13.3 mm^2^. Referring to Fig.(1), we chose a reference where we indicated: with x the beam propagation direction; with y the pivoting direction and the light-sheet width; with z the detection direction and the light-sheet thickness.

**Figure 1.**
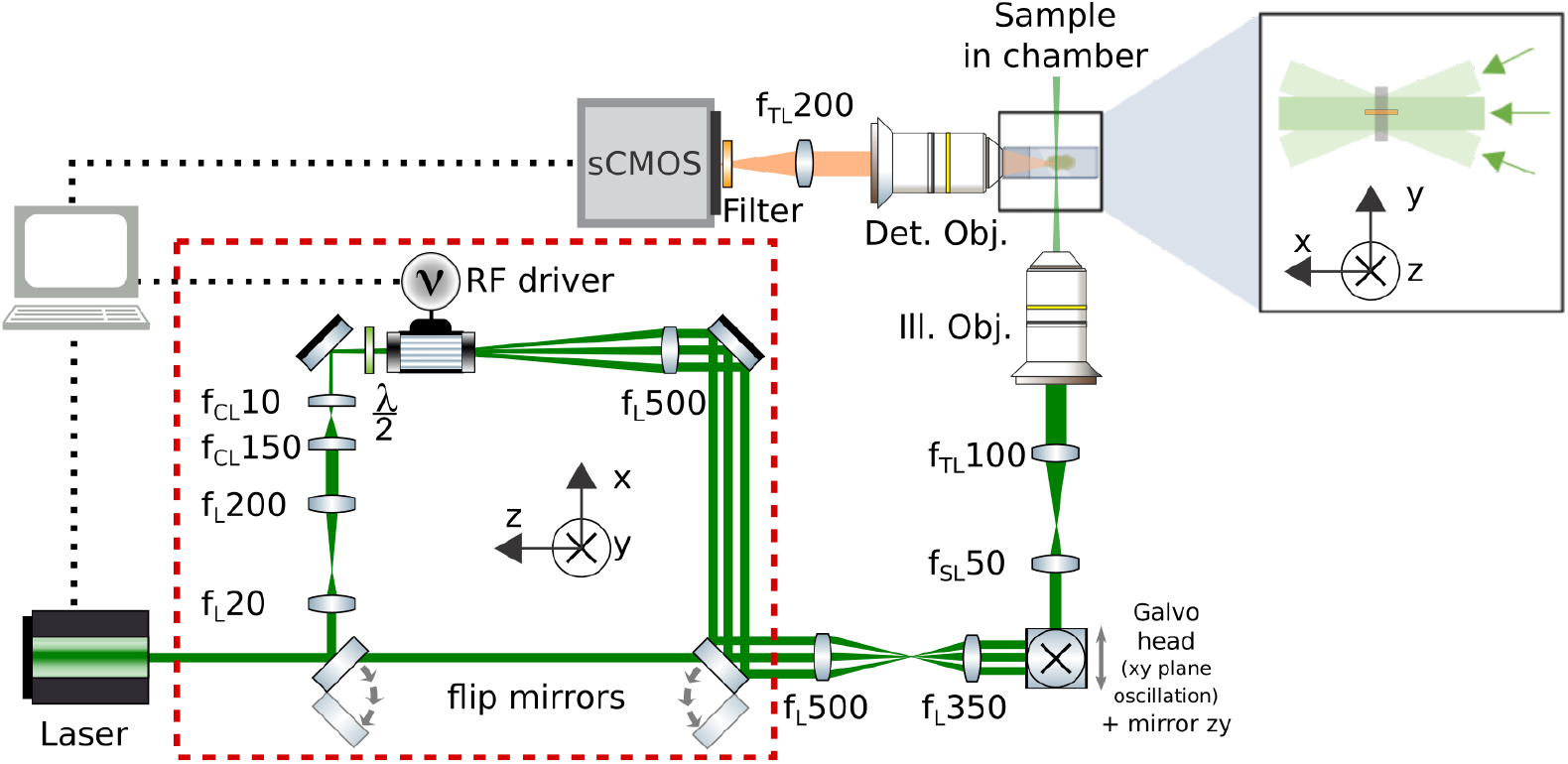
Schematic of the multi-directional DSLM setup, presenting the excitation and imaging paths. The red dashed rectangle denotes an alternative light path, selectable via flip-flop mirrors, where two cylindrical lenses CL shape the circular Gaussian beam into an elliptical profile that is then pivoted by an AOD.

The experiments were carried out in the confocal detection regime, obtained by defining a digital slit through a row of pixels simultaneously activated on the camera sensor (the dimension of such slit was of 221 μm in camera space). To verify that confocal detection effectively improves the image contrast, we performed acquisitions also in the widefield regime by leveraging the global shutter camera modality. Specifically, we maintained the same line exposure time to keep constant the per-row illumination intensity, while the digital slit height was expanded by about ten times via a corresponding reduction of the rolling shutter sweep pace (up to 2048 μm in camera space). The detection parameters are reported in the following table 1 and were different between the PSF measurements, using fluorescent beads, and the mouse brain imaging.

**Table 1.** Detection parameters used for PSF measurements with fluorescent beads and during mouse brain imaging.

| Experiment | N. frames | Step size | Line exposure time | Rolling shutter pace |
| --- | --- | --- | --- | --- |
| PSF est. (confocal det.) | 200 | $1 \mu\text{m}$ | 3 ms | $90.00 \mu\text{s}$ |
| Brain imaging (confocal det.) | 350 | $2 \mu\text{m}$ | 3 ms | $90.00 \mu\text{s}$ |
| Brain imaging (widefield det.) | 350 | $2 \mu\text{m}$ | 3 ms | $9.74 \mu\text{s}$ |

### 2.2. Animal experimental procedures

Male FosTRAP mice (B6.129(Cg)-Fos^*tm*1.1(*cre*/*ERT* 2)*Luo*^/J x B6.Cg-Gt(ROSA)26Sor^*tm*9(*C AG*−*tdTomato*)*Hze*^ /J, n=3) were used for this work. Adult mice were handled and injected with saline solution daily for at least 3 days prior the 4-hydroxytamoxifen (4-TMX) injection, always leaving them in their own homecage. 4-TMX (Sigma H6278) was first dissolved in ethanol to a concentration of 20 mg mL^−1^. This stock was then melted with corn oil at 37 ◦C to obtain an injectable oil formulation. The last day, 50 mg kg^−1^ of 4-TMX was given intraperitoneally to all mice. All experimental procedures were approved by the Italian Ministry of Health (Authorization n. 512-2018_FC) [37].

### 2.3. Ex-vivo processing and CLARITY

One week after the tamoxifen injection, animals were deeply anesthetized and transcardially perfused with 50 mL of ice-cold 0.01 *M* phosphate buffered saline (PBS) solution (*pH* 7.6) followed by 75 mL of freshly prepared paraformaldehyde (PFA) 4 % (*pH* 7.6). The extracted brains were then processed according to the CLARITY/TDE protocol [38]. In detail, specimens were left in PFA at 4 ◦C over night. The following day, samples were incubated in the hydrogel solutions (containing 10 % acrylamide (wt/vol), 2.5 % bis-acrylamide (wt/vol) and 0.25 % VA044 (wt/vol) in PBS) at 4 ◦C for 3 days to allow a sufficient diffusion into the tissue. Samples were then degassed, replacing oxygen inside the vials with nitrogen, and the hydrogel was polymerized by incubation in water bath at 37 ◦C for 3 hours. Later, embedded brains were then placed in clearing solution (full of sodium dodecyl sulfate, SDS) at 37 ◦C. Specimens were gently shaken throughout the whole period. After clearing, samples were incubated 1 day in PBS with 0.1 Triton-X (*pH* 7.6) and 1 day in PBS (*pH* 7.6), removing the excess SDS. Finally, murine brains were immersed in a mixture of 40 % 2-2’ Thiodiethanol (TDE) in PBS (vol/vol) for imaging, that has a refractive index *n*_refr_ ≈ 1.41.

## 3. Results

To evaluate the light-sheet pivoting effect on striping artifacts, we implemented an optical set-up with two alternative paths for the excitation laser beam, conveniently selectable through flip mirrors. Briefly, through the first optical path, the sample is illuminated with a standard pencil-like Gaussian beam, as used in a DSLM (this configuration is labelled with NO AOD in the rest of the paper). In the second configuration, the path differs mainly by the presence of an AOD and two cylindrical lenses which collapse one dimension of the circular Gaussian beam into an elliptical profile, creating a “mini light-sheet”. Inserting the AOD in the optical path before the galvanometric scanning head allows to implement beam pivoting. In practice, the field of view containing the sample is illuminated line by line by the galvo head, while the AOD rotates the excitation beam around its propagation axis at a much faster rate.

We compare the results provided by the first mentioned configuration with two alternative illumination approaches allowed by the layout with the AOD. The first is a multi-beam arrangement of a user-selectable number of mini light-sheets that propagate simultaneously at different angles (in sequence one, three and five static beams, respectively denoted as 1LS, 3LS, 5LS), while the second is a single mini light-sheet dynamically pivoted over the entire angular range and labeled as scanning beam (SB).

### 3.1. Experimental set-up characterization

We first calibrated the AOD used to tilt the beam. In such a device, a radio-frequency (RF) is applied to a piezotransducer to generate a pressure wave that propagates through the device’s internal crystal, acting globally as a diffraction grating for the beam with fine control over its deflection angle and intensity. If the piezotransducer is driven by multiple frequencies, then a linear combination of gratings is produced, allowing to generate simultaneously different beams from a single one, with independent regulation in terms of spatial direction and intensity. To evaluate the angular deflection as a function of the driving radio frequency, a single beam was projected inside the sample chamber filled by a 40 % 2-2’ Thiodiethanol solution in water, without the sample. The input RF frequency signal was changed continuously, acquiring three images for each setting, and the corresponding tilt angle was measured using the angle tool of the open-source software ImageJ. We selected as central frequency the one with which the beam entered into the sample chamber perfectly parallel to the optical axis, i.e. the one orthogonal to the detection direction. Then a frequency sweep spanning [−5; +5] MHz was defined around this value to cover a large enough pivoting angle to envelope the digital slit used in confocal detection. Figure (2A) shows the measured AOD’s angular response as a function of the relative shift from the central frequency. A linear model has been fitted to the data, with a R-square parameter of 0.9989, finding an angular coefficient *m* = 1.25 ± 0.03° MHz^−1^ between the deviation angle and the frequency shift.

**Figure 2.**
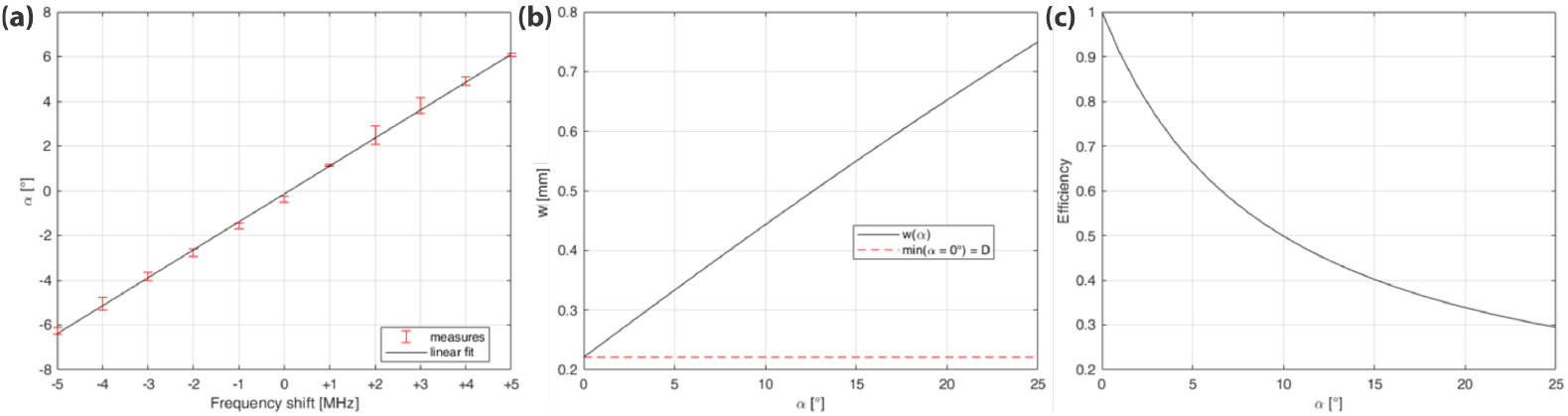
Setup characterization: (A) shows the AOD’s angular response as a function of the input radio frequency shift relative to the referenced central value. (B) and (C) show the simulated dependence, respectively, of the beam width w and of the detection efficiency *E f f* from the pivoting angle *α*, up to the maximum angle admitted by the excitation objective.

Confocal detection on a sCMOS sensor is realized using the rolling shutter modality. A digital slit is made by simultaneously active pixel rows that roll through the pixel matrix synchronously and aligned with the illumination beam, thus improving the image contrast and rejecting out-of-focus fluorescence light coming from the sample. The digital slit has finite dimensions in sample space, with width and height defined respectively by the horizontal FOV and by the virtual slit width on the camera, which is set automatically when the rolling shutter speed and the line exposure time are chosen by the user. Fig.(3) shows a schematic of the beam pivoting geometry, as observed during confocal detection at the digital slit on the sCMOS camera. As shown, the beam is tilted inside the sample chamber, where *w* is the beam width, *D* is the virtual slit width, *L* is the length of the digital slit corresponding to the camera FOV and *α* is the pivoting angle which defines also the complementary angles *β* + *δ* = *π* / 2 − *α*.

**Figure 3.**
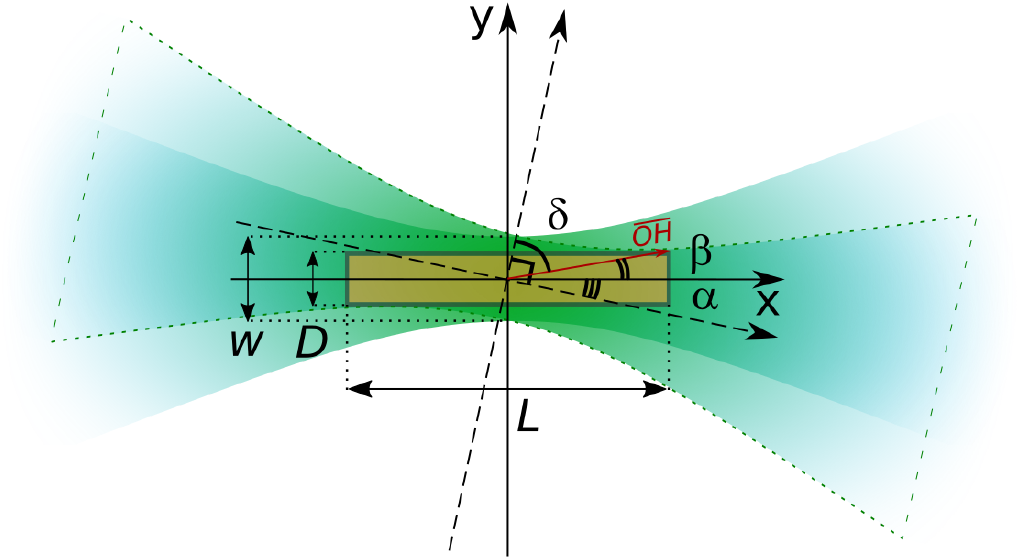
Schematic of the beam pivoting arrangement at the sample: *w* is the beam width; *D* is the virtual slit width; *L* is the length of the digital slit on the camera sensor (sepia rectangle); α is the pivoting angle which defines also the complementary angles *β* + *δ* = *π*/2 − α.

Pivoting a pencil-like Gaussian beam would make large parts of the beam rotate outside the physical dimensions of the digital slit, leading to uneven illumination. This problem can be solved by optically shaping a circular Gaussian beam into an expanded elliptical-Gaussian beam with its width tuned to cover the whole aperture at each inclination. In the following, we introduce the optical model that enabled us to engineer the second optical pathway such that it fulfills this requirement.

The semi-diagonal of the slit, indicated with 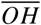, can be easily calculated as:

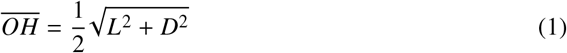

The beam half-width *w*/2 can be found from

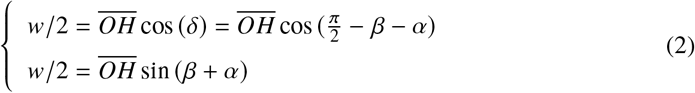

while *β* is

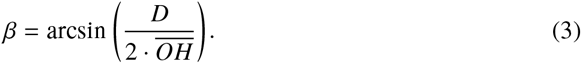

That means there is a strict relation between the maximum angle *α* used to pivot the beam and the beam width w. The detection efficiency can be accounted for by considering the difference between the effective detection area, indicated with the small sepia rectangle in Fig.(3), and the area illuminated when scanning the beam through the sample, denoted by the green rays. We can define this parameter simply as the ratio between the two corresponding dimensions:

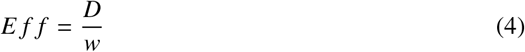

Figures (2B) and (2C) show the dependency of these parameters on the pivoting angle *α*.

The digital slit dimension was *D* = 221 μm in camera space (see Section 2 for further details) and the length of the digital slit was *L* = 1.3 mm, from which we found 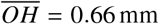. The frequency range set on the AOD to scan the beam has been fixed at [68 ÷ 72] MHz around the central value of 70 MHz, covering a total angular range of Δ*θ* = 2*α* = 5°. The RF power for each applied frequency was set to 16 dBm and 25 dBm, respectively in the multi-beam and SB modes. Consequently, the beam half width required to fulfill the demands of pivoted confocal illumination is then *w*/2 ≃ 140 μm. This leads to an expected detection efficiency of *E f f* = 79 %.

The optical path containing the AOD and the cylindrical lenses has been developed according to these considerations. Our beam-shaping system produces an elliptical Gaussian beam profile with numerical apertures NA_*z*_ = 0.022 and NA_*y*_ = 0.0015. The resulting waist along the detection direction is *w*_*z*_/2 = 8 μm, which leads to a Rayleigh length of the virtual light-sheet of 530 μm along the illumination propagation direction, providing, thus, an almost uniform illumination over the FOV. The waist along the pivoting direction is *w*_*y*_/2 = 122 μm, which is very close to the desired optimum value and provides for a small beam divergence, differently from the configuration presented in [34]. The high degree of angular diversity required for efficient shadowing suppression is brought by beam pivoting, that results in an effective numerical aperture of 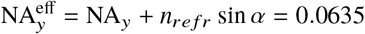.

### 3.2. PSF Estimation

To quantify the characteristics of the light-sheet microscope, the PSF has been measured for both optical path configurations, the one with AOD and the second one without. We prepared a specimen containing fluorescent beads included in 4 % agarose gel. TetraSpeck™ microspheres (Invitrogen T7279, with radius *r* = 50 nm) were used at a final concentration of 0.0025 % (vol/vol). The intensity profiles along the radial and axial directions of n=6 sub-micrometric fluorescent beads have been computed using the open-source software ImageJ. Each has been fitted with a Gaussian model to calculate its full width at half maximum (FWHM). Fig.(4) shows the raw data points and the Gaussian model computed with the mean FWHM, obtained respectively along the radial (panels (a)-(c)) and axial (panels (d)-(f)) directions. In detail, from left to right one, can observe the results obtained respectively for the NO AOD, 3LS and SB configurations. The two panels (g) and (h) show the mean and the standard deviation of the FWHM extracted from these measurements. The mean FWHM is found to be approximately three times larger than the theoretical lateral resolution, most probably due to aberrations caused by refractive index mismatch in the detection path. As one can observe, the alternative optical path, where the AOD is introduced to pivot the beam, does not affect significantly the lateral and axial resolution. Panel (i) shows a representative frame containing the fluorescence signal produced by a fluorescent bead.

**Figure 4.**
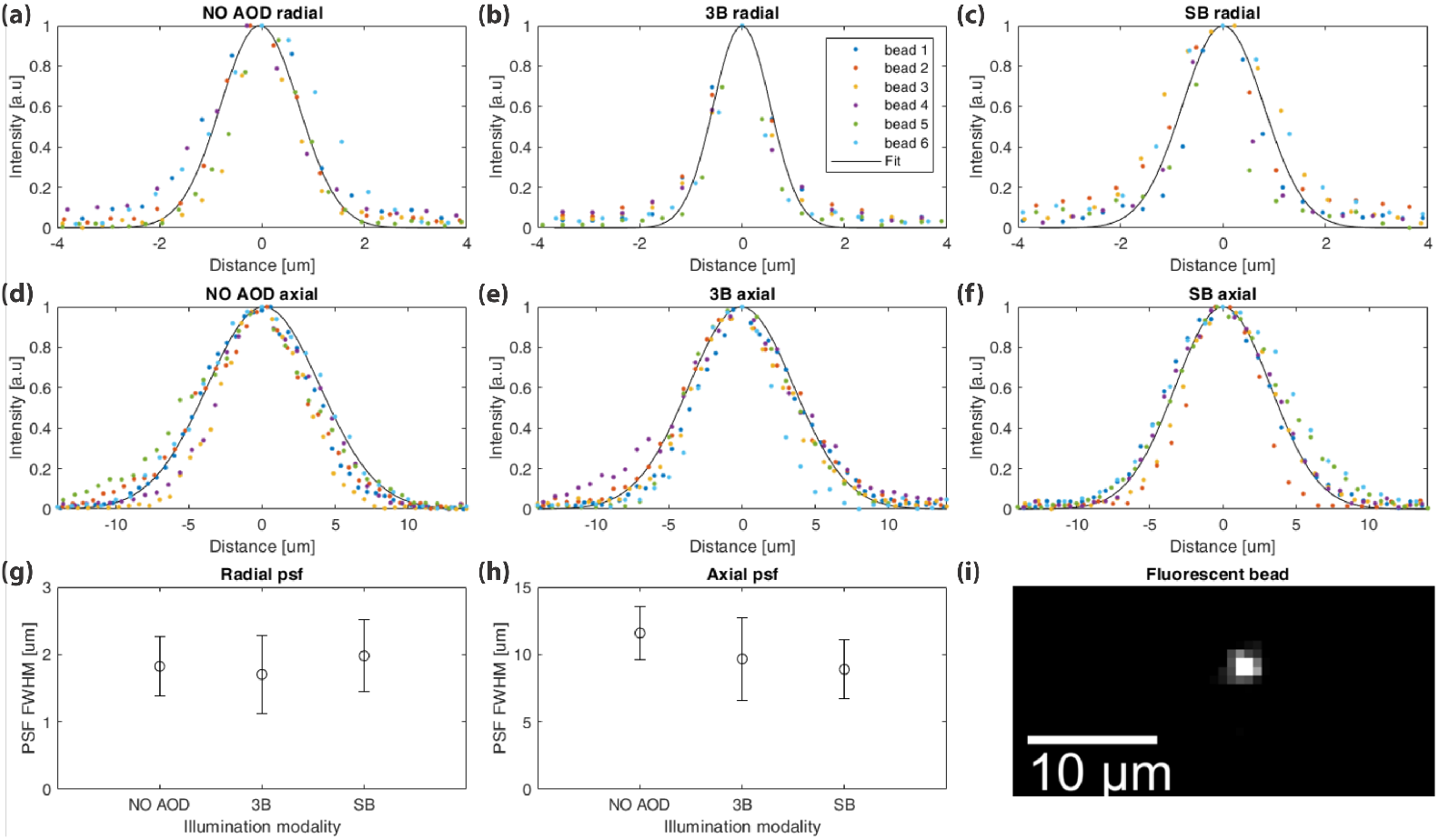
Panels (a)-(c) and (d)-(f)) show the raw data points (encoded by a different color for each of the 6 measurements) and the average Gaussian fit (black line), obtained respectively in radial and axial direction. From left to right, the results obtained respectively for the NO AOD, 3LS and SB configurations. Panels (g) and (h) show the average FWHM and the standard deviation extracted from these measurements. Panel (i) shows a representative frame containing the fluorescence signal detected from a Tetraspeck fluorescent microsphere (*r* = 50 nm) embedded in agarose gel. Scale bar size: 10 μm.

### 3.3. Imaging of mouse sample

To prove the advantages of the presented illumination approach on biological tissues, we imaged with all configurations of our multi-directional DSLM three cleared mouse brains, expressing the fluorescent protein tdTomato (details regarding sample mounting and preparation in Sec.2). The right column in Fig.(5) shows the same view in each modality, while the left reports the normalized gray-scale intensity profiles taken along the vertical dotted red arrow. There is no evident difference between the classic NO AOD configuration and the 1LS with just a single mini light-sheet created by the AOD. This was expected because they are optically equivalent. To solve the striping problem, one needs to dynamically tilt the light-sheet to average out the shadows attenuation or, alternatively, have more than one mini-light-sheet simultaneously present through the sample. Indeed, from the 3LS up to 5LS configurations the shadow artifacts are gradually reduced, as well as in the SB mode. In particular, the two latter approaches present a very similar performance.

**Figure 5.**
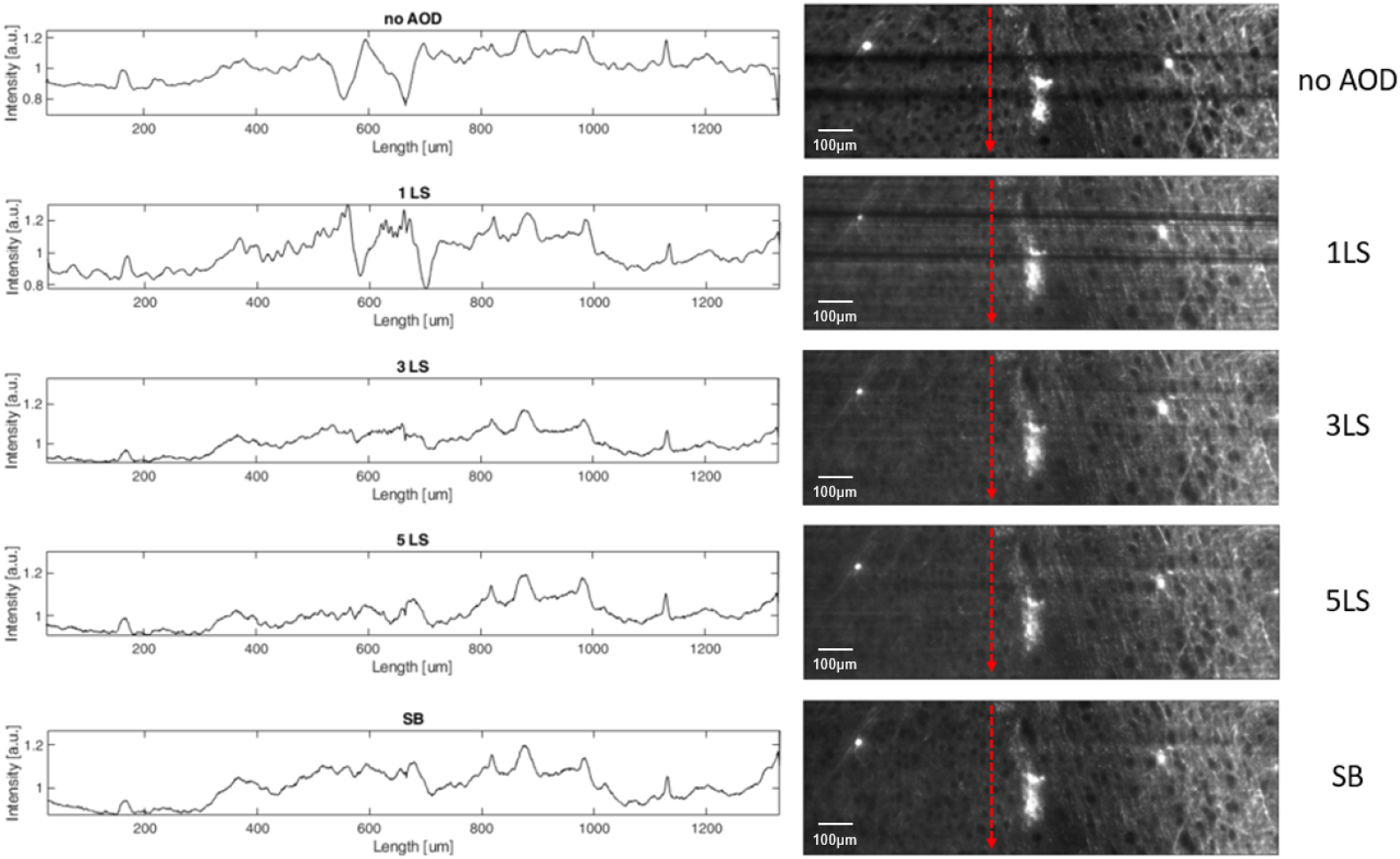
On the right LSFM images of a mouse brain, expressing fluorescent protein tdTomato, obtained by pivoting the beam with different scanning configurations, respectively from up to down, NO AOD, 1LS, 3LS, 5Ls and SB. On the left, the normalized gray-scale intensity profiles taken along the vertical directions. Scale bar size: 100 μm.

To quantify the shadowing suppression, we analysed the stacks acquired in SB and NO AOD configurations and belonging to different areas of three imaged mouse brains. The left panels of Fig.(6) show the normalized intensity profiles of the frames displayed on the right, each obtained from a longitudinally averaged intensity projection, together with their difference. The degree of shadowing suppression can be estimated from the ratio *R* between the integrals of the absolute values of the difference intensity profile and the SB one, taken as reference. We considered substacks of ten consecutive frames, on which we calculated the mean and standard deviation of the ratio. The results reported in Table 2 show an effective improvement provided by pivoting the beam.

**Figure 6.**
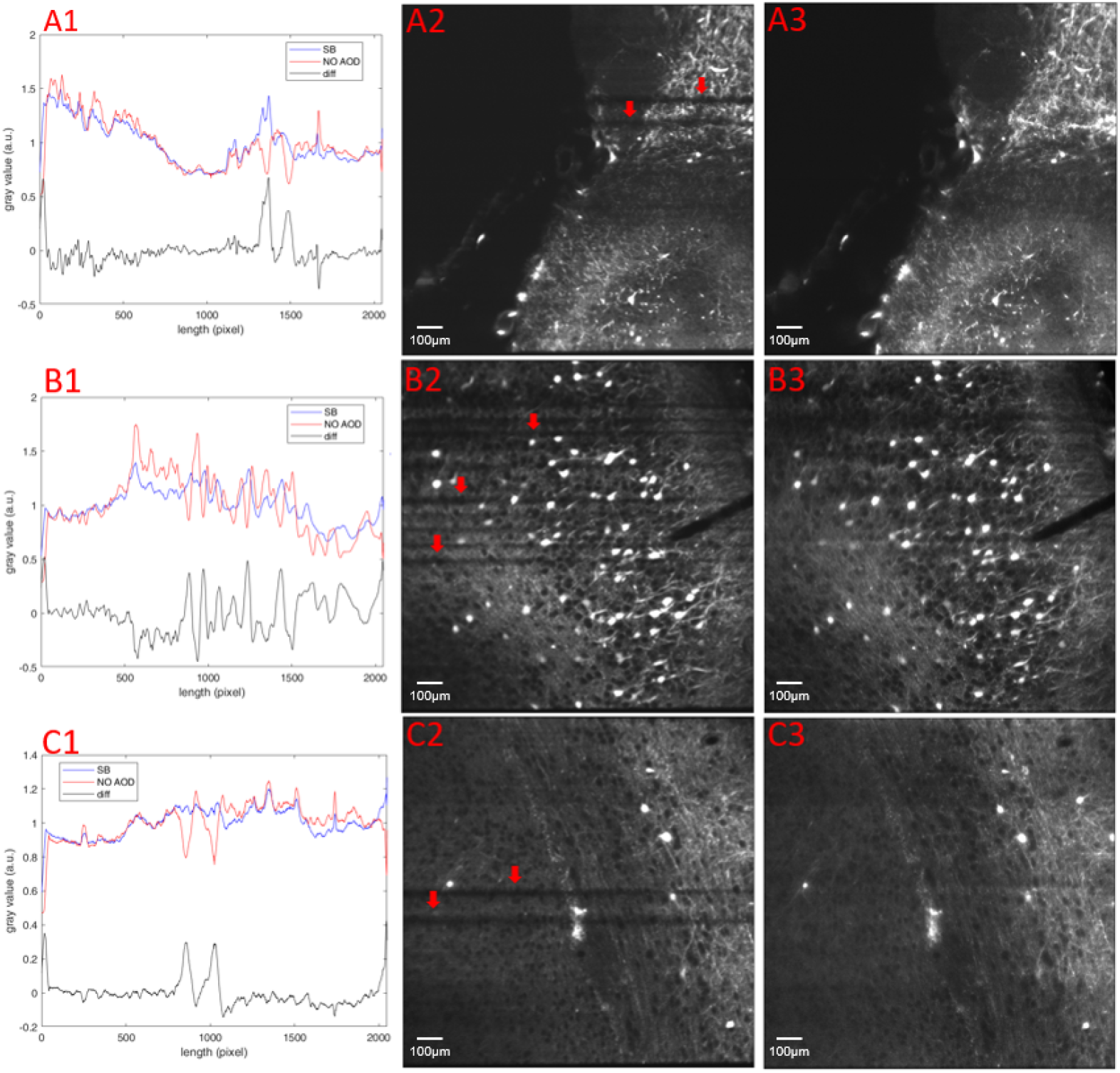
Shadowing suppression quantification. (A1), (B1) and (C1) show the normalized intensity profiles, together with their corresponding difference, obtained by a longitudinally averaged intensity projection of the images; (A2), (B2) and (C2) show single frames acquired in three different mouse brains expressing fluorescent protein tdTomato, taken in NO AOD configuration; correspondingly, A(3), B(3) and C(3) represent the same view taken with SB scanning configuration. Areas of larger striping attenuation are indicated by red arrows. Scale bar size: 100 μm.

**Table 2.**
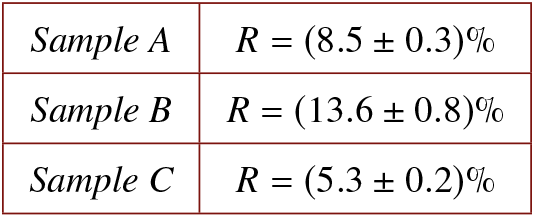
Shadowing suppression ratios, obtained by a longitudinally averaged intensity projection of a substack of ten consecutive frames.

|  |  |
| --- | --- |
| *Sample A* | $R = (8.5 \pm 0.3)\%$ |
| *Sample B* | $R = (13.6 \pm 0.8)\%$ |
| *Sample C* | $R = (5.3 \pm 0.2)\%$ |

### 3.4. Contrast analysis

Confocal light-sheet microscopy allows to reduce out-of-focus contributions from scattered photons, improving the image contrast with respect to widefield detection in the global shutter acquisition mode, where all camera pixels are active simultaneously. To test and quantify such improvement and evaluate the eventual impact of pivoting on it, the image contrast ratio between these two configurations has been calculated for the SB and NO AOD illumination modalities, according to the normalized discrete cosine transform (DCT) Shannon entropy [7]. In table 3, we indicated with *CR*_SB_ and *CR*_NO AOD_ respectively: the ratio between the contrast calculated on images taken with SB configuration with the digital slit and the one obtained on images without slit; the ratio between the contrast obtained using NO AOD configuration, with the digital slit and without. The data was taken from several brain areas in three different samples. The values refer to the mean and standard deviation calculated from all the contrasts which are evaluated by comparing each corresponding frame between the acquired stacks. The reported data indicates a clear and statistically significant improvement in contrast for both modalities, demonstrating the benefit of confocal detection. The third column displays the P-value calculated between the two data sets. The contrast ratio enhancement with illumination pivoting is smaller than without and this difference may be attributed to the not perfect confocality linked to pivoting. The beam can indeed excite some areas out of the digital slit, while being swept for shadowing reduction.

**Table 3.**
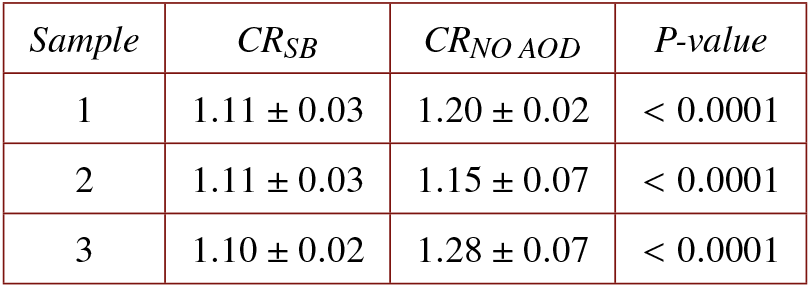
Contrast enhancement by confocal detection, calculated over stacks acquired with SB and NO AOD configurations.

## 4. Discussion

Light-sheet Fluorescence Microscopy is a widespread technique in neuroscience for 3D imaging of extended biological samples and small animal whole brain anatomy reconstruction. However, images acquired with LSFM are often affected by striping artifacts, typically caused by shadows originated from absorbing or scattering objects inside the sample that partially block the illumination.

Here we demonstrated an optical solution that allows to concurrently attenuate the striping artifacts and preserve the contrast enhancement provided by confocal detection, with an almost uniform FOV illumination. It is based on generating a fast multi-directional illumination of the sample via an AOD that controls a tailored elliptical-Gaussian beam, provided in turn by a simple optical beam-shaping system. The illumination numerical aperture resulting from such approach is consequently anisotropic with NA_*z*_ = 0.022 and NA_*y*_ = 0.0015. The subsequent beam pivoting leads to an effective increase in the numerical aperture 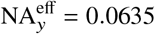. Striping artifact suppression is deeply correlated with the illumination numerical aperture because the size of the occluding objects defines the smallest incidence angles required for the light to circumvent them, while the latter sets the angular diversity of the impinging beam. Taking advantage of large numerical apertures, as in [34], intrinsically provides enough angular diversity to result in shadowing attenuation, recovering information behind the obstructions by looking from several angles. However, this reduces the illumination uniformity over the FOV and requires image tiling for optimal results [35]. Another route to increase the effective numerical aperture, while avoiding tiling, is to pivot a small NA beam around its own propagation axis.

With the AOD we are able to pivot the beam with respect to the propagation axis, with a scanning rate faster than the detector acquisition rate, to average out over time the shadowing attenuation from different angles. Moreover, with AODs, multiple beams can be generated and projected at diverse angles simultaneously: in such a way we implemented several scanning configurations without imposing any constraint on the imaging rate. Realizing such flexible pivoting illumination scheme with galvo mirrors would require a more complex optical system, while remaining potentially limited in the peak sweeping rate by their intrinsic mechanical inertia.

Pivoting represents also an alternative solution with respect to non-Gaussian beam illumination, like Bessel beams. The applicability to LSFM of such beams has been demonstrated together with their capability of shadow suppression, even if they show reduced image contrast and signal-to-noise ratio with respect to Gaussian beam illumination. On the other hand, the Bessel beam light distribution is an issue for sample photo-bleaching and photo-toxicity, carrying out-of-focus contributions to the illumination. Such problem is still present when one wants to adapt the pivoting approach to CLSFM because a large part of the rotating illumination beam would not be collected by the camera sensor. In this regard, we found a fully optical solution by modeling the illumination and pivoting geometry, determining, thus, the beam shape requirements to fill the digital aperture for all angles. Using this information, we built a simple optical beam-shaping system that produces the recommended tailored elliptical-Gaussian illumination beam.

The PSF response has been measured for two illumination configurations, one with a standard pencil-like Gaussian beam, as used in DSLM, and, alternatively, one which exploits the AOD pivoting features. It has been shown that the second optical path does not affect significantly the lateral and axial resolution while pivoting.

In order to observe the advantages carried by the AOD pivoting in terms of shadow suppression, we imaged three cleared mouse brains expressing the tdTomato fluorescent protein, showing an effective improvement with respect to the classical approach, both in the scanning beam and multi-beam approaches.

We finally verified the image contrast enhancement provided by confocal detection with respect to widefield imaging (in the camera global shutter acquisition mode) for both the classical Gaussian beam illumination and the one with the pivoted beam. However, the contrast improvement was found to be larger for the classical approach, because in the pivoting configuration part of the illumination spreads out from the slit. Additionally, dynamically sweeping the beam around the propagation axis results in a lower average illumination power over the excited volume with respect to static illumination approaches, requiring, thus, a higher laser excitation power to compensate.

## 5. Conclusions

We demonstrated that the presented scanning AOD system is compatible with digital scanned laser light-sheet fluorescence microscopy. We tested its performance by acquiring several mouse brain areas, observing real-time shadow suppression, while preserving the contrast enhancement of confocal detection by using a pivoted elliptical-Gaussian beam.

Such fast and flexible confocal DSLM with reduced striping artifacts may benefit high throughput imaging of large tissue volumes [1, 5, 15, 27] and live functional studies with high temporal resolution [8, 39, 40]. Due to the widespread use of LSFM in live imaging experiments, e.g. fast processes monitoring or biological dynamic tracking [10–14], a further check upon our illumination approach suitability for live biological samples would be of interest. Recovering spatio-temporal information lost to striping artifacts during neuronal activity recording, for example in zebrafish larvae in specific pathological or physiological conditions [25,26,31], would be a major step forward in neuro-imaging. Due to the larger laser excitation power required by this approach with respect to classical scanning modes, particular attention should be paid to sample photo-toxicity, especially in live imaging.

